# Extreme fatigue-resistance in a brood parasite parallels a war of attrition in begging competition with hosts

**DOI:** 10.1101/2024.10.31.621188

**Authors:** Nicholas D. Antonson, Mark E. Hauber, Wendy M. Schelsky, Carly Kallembach, Matthew J. Fuxjager

**Affiliations:** Department of Ecology, Evolution, and Organismal Biology, Brown University, Providence, RI, USA; Department of Evolution, Ecology, and Behavior, University of Illinois at Urbana-Champaign, Urbana, IL, USA; Advanced Science Research Center and Program in Psychology, Graduate Center of the City University of New York, New York, NY, USA; Illinois Natural History Survey, Prairie Research Institute, University of Illinois at Urbana-Champaign, Champaign, IL, USA

**Keywords:** begging behavior, brood parasitism, *musculus complexus*, muscular fatigue, host-parasite coevolution, physiological constraints

## Abstract

Food-solicitation displays are critical to the survival of dependent young. These signals often evolve in extreme ways in response to familial conflict among siblings and between parents and offspring. Avian brood parasites demonstrate how these signals can become exaggerated by engaging in a begging arms race with host offspring. However, despite hypotheses about how begging displays in brood parasites are adapted to support the demands of early parasitic life, we know little about the physiological specializations that support exaggerated parasitic traits. Here, we investigate display performance in the begging competition between brood parasitic brown-headed cowbirds (*Molothrus ater*) and host prothonotary warblers (*Protonotaria citrea*). Using *in situ* muscle stimulation, we test the performance limits of the musculus complexus (**MC**), which actuates begging displays. Our results show that nestlings of each species initiate their begging display at similar speeds, but cowbirds beg far longer than warblers. We trace these effects to species differences in muscle performance, with the cowbird **MC** showing dramatically greater fatigue-resistance than warblers, while other facets of muscle performance remained similar between species. Altogether, these findings suggest that intra-brood conflict in host-parasite interactions selects for extreme fatigue-resistance in skeletal muscles to support a “war of attrition” in begging behavior and nestling survival.

## Introduction

In multiparous species, the allocation of food by parents generates evolutionary conflict between offspring [1]. This is because food is often a limited resource, and parents must optimize its allocation to maximize their own lifetime fitness outcomes. On the other hand, offspring must prioritize their own individual interests over those of parents or siblings [2,3]. Such misalignment of interests can generate strong selection for elaborate traits that favor the evolution of sibling competition through signaling traits such as ornaments and food-solicitation displays [4–6]. Thus, intense conflicts are often matched with increasingly selfish and exaggerated food-solicitation signals to compel parents to feed a specific offspring [1]. Theoretically, trait innovation to exaggerate such displays should occur through the concomitant evolution of supportive physiological mechanisms [7]; yet, few studies have ever explored such mechanisms, or how they underlie the expression of selfish behaviors.

Here, we study food-solicitation in nestling birds, which takes the form of begging displays: chicks raising their heads, opening their beaks, and calling to parents. Studies have not only shown that such displays are the first coordinated behavior that birds perform post-hatching, but also that these displays are critical for survival [7]. When begging, nestlings engage their neck muscles to throw back their head, extend their neck, and expose their mouths to provisioning parents. By simply maintaining the postural height of their begging display and position in the nest for longer durations, chicks can greatly increase their provisioning success, and hence their growth, survival, and recruitment [6,8–10]. Therefore, superiority in the begging arms race amongst siblings should favor the evolution of mechanisms that support the exaggeration of these gestural traits if not curbed by parental neglect or other contrasting selective pressures [11].

Nowhere is this begging arms race more apparent than in obligate brood parasites, where one species lays its parasitic egg in the nest of a host species. Once parasitic chicks hatch, they are often in direct competition with host offspring for parental resources. In the resulting coevolutionary arms race, parasitic chicks are genetically unconstrained from any inclusive fitness mechanisms that could regulate how selfish phenotypes evolve amongst true siblings [12,13]. Thus, parasitic chicks are free to evolve strategies to usurp unrelated nestmates, such as highly exaggerated begging displays that help them bias provisioning of food from host parents in their favor [14–17].

Here, we study the physiological basis of a well-known case of exaggerated begging [14] in generalist brown-headed cowbirds (*Molothrus ater*), and we compare the display to that of one of the species’ many hosts, prothonotary warblers (*Protonotaria citrea*). Specifically, we measure performance attributes of the *musculus complexus* (**MC**), a large, striated neck muscle [18] that actuates dorsal flexion and extension of the neck for both pipping through eggshells during hatching and begging displays as nestlings [19,20] (**Fig 1A**). The **MC** is primarily composed of singly-innervated fast-twitch fibers and few slow twitch fibers, suggesting that selection drives nestlings to assume begging postures quickly, but with little endurance [7]. Accordingly, we hypothesize that parasitic cowbirds out-beg host nestlings through a war of attrition, whereby both species respond to a provisioning stimulus at similar speeds, but only cowbirds can maintain this posture for extended periods of time to help exaggerate their display. Further, we hypothesize that the underlying **MC** of parasitic cowbirds and host warbler nestlings mirrors this behavioral asymmetry when subject to an *in situ* performance assay designed to probe muscular resistance to fatigue.

**Figure 1.**
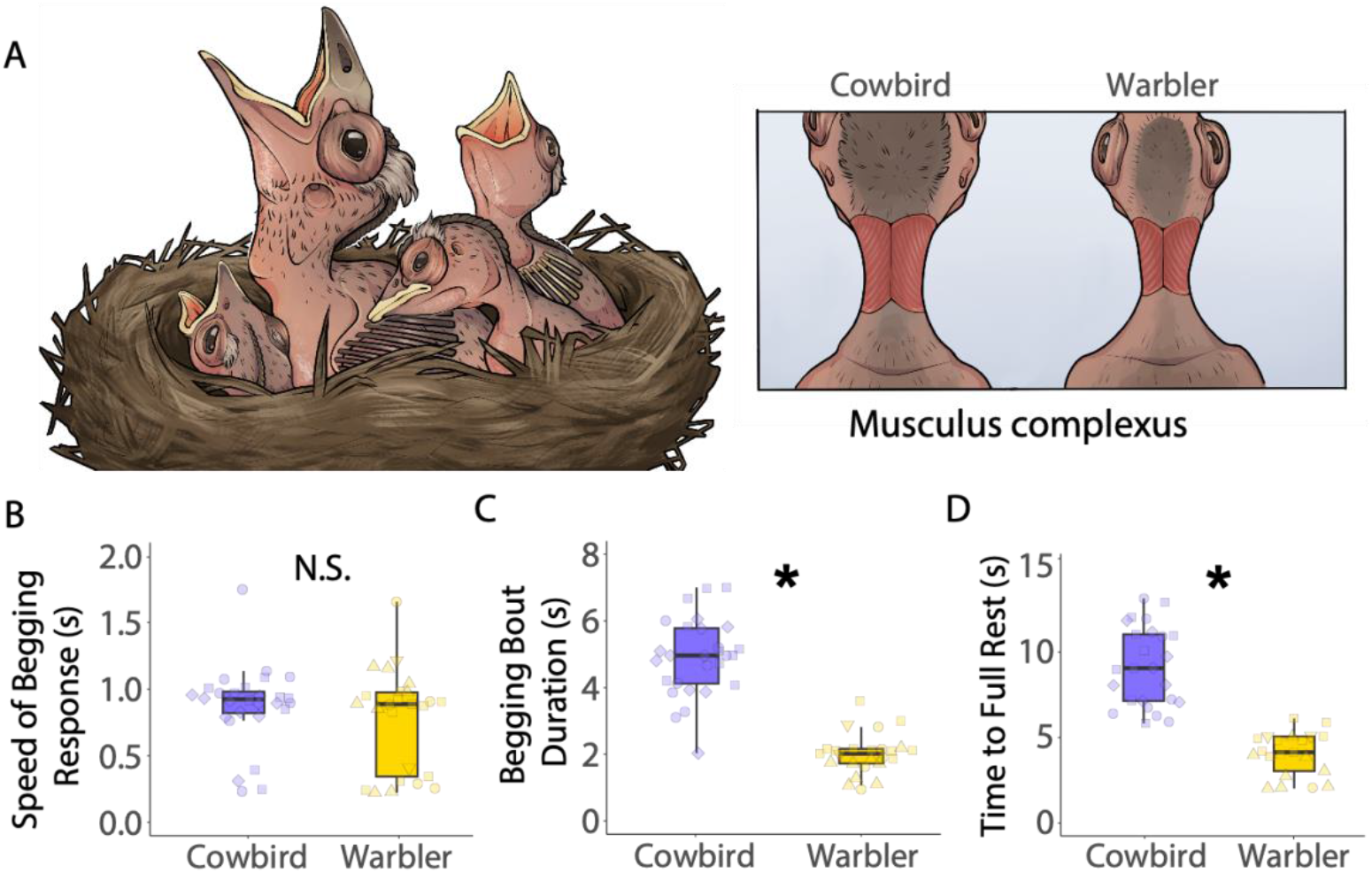
Hunger-driven begging performances of cowbirds and hosts. A) Illustration by CF Stowers of a cowbird nestling in begging competition with three host warbler nestlings and a schematic of the *musculus complexus* (**MC**). B) Speed at which cowbirds and warblers assumed peak begging posture in response to acoustic stimulus was statistically indistinguishable (N.S., p>0.05). C) Cowbirds begged with the neck extended for significantly longer than warblers. D) Cowbirds spent more time performing begging-associated activities than warblers. Box plots show interquartile-range and median. Asterisk (*) denotes significance (p<0.05), and shapes represent n=3 cowbirds and n=4 warblers, with number of each shape showing replicates.

## Materials and Methods

### 1. Study system

This study took place in artificial nest boxes occupied by prothonotary warblers (*Protonotaria citrea*) in the Cache River watershed of southern Illinois, USA in 2021. Prothonotary warblers are neotropical migratory songbirds and obligate secondary cavity nesters that preferentially nest in cavities above standing water. Their nests are regularly parasitized by the generalist obligate brood parasitic brown-headed cowbirds (*Molothrus ater*) [21]. Prothonotary warblers readily breed in artificial nest boxes, often preferring them over natural cavities [22]. Nest boxes were designed and protected from predators and ectoparasites and nests were monitored daily from clutch initiation through hatching to ensure accurate age assessment as in Antonson et al. [23].

Cowbirds most often experience 1 day of hatch asynchrony ahead of their hosts, so all experimental procedures were carried out on warblers that were 1 day younger than the cowbirds studied to mirror ecologically-relevant asymmetry in developmental stage [24,25]. A comprehensive meta-analysis on host hatching asynchrony is not currently published. However, we collated data suggesting that only 36 species of hosts hatch at the same time or prior to cowbirds, while 176 species lag behind cowbirds in development (Antonson unpublished data). Further, evidence exists that cowbirds adjust their growth rates in response to variation in hatch asynchrony across different host species [26]. In this regard, prothonotary warblers typically hatch a day after cowbirds and remain in the nest for slightly less than 11 days before fledging while cowbirds fledge at approximately 11 days in the population studied [27], making them an ideal host for testing hypotheses on physiological performance that may be contingent on developmental stage. Brood sizes contained 3 or 4 warblers with a cowbird for these experiments. The largest warbler in each nest was used for focal begging and physiology comparisons with the cowbird. All procedures addressed here were approved by the University of Illinois at Urbana-Champaign Institutional Animal Care and Use Committee.

### 2. Hunger-driven begging assays

On Day 5 after hatching for warblers (n=4), and Day 6 after hatching for cowbirds nestlings (n=3), chicks were removed from the nest box and placed in an externally warmed, dark nest box on the ground. Technical difficulties with video equipment prevented data collection from a 4^th^ cowbird. The cowbird and warbler from a focal nest were removed at the same time to ensure removal did not cause variation in provisioning that may affect hunger-driven begging performances. The begging assay nest boxes contained a mix of soft nesting material and moss shaped into a nest cup. An infrared camera attached to a digital video recorder (DVR) was threaded into the nestbox to record video. Nestlings were observed in silence for 30 minutes to induce hunger during which time researchers observed virtually no spontaneous begging. At 30 minutes, researchers tapped continuously on the side of the nestbox using a #2 pencil to stimulate begging. Tapping continued until after the bird returned to rest and occurred 5 times in a row with 15-minute breaks in between rounds of the assay. As this was a test of how long the birds could keep there heads up in a continuous and discrete bout, rest was defined as the first point at which the nestling rested its head against the side of the nest box or the nest substrate. We chose this method as any amount of rest would allow a degree of muscle recovery that would have been inconsistent with our later *in situ* muscle performance assays. Additionally, this discrete change in behavior (return to rest) was chosen because it represents a conservative threshold that is unlikely to be affected by observer bias. At 1 hour out of the nest, nestlings were removed from the assay box, fed fresh-mixed baby bird formula to satiation, and returned to the focal nest.

### 3. In situ muscle stimulation

#### 3.1 Experimental Setup

To evaluate the fatigue-resistance and twitch kinetic properties of the **MC**, we used an *in situ* stimulation approach that has been implemented in a range of species, including humans [28– 32]. *In situ* stimulations showcase muscle performance under natural load and length conditions, where the muscle receives *in vivo* delivery of oxygen and energy to power movement. On Day 6 for warblers (n=4) and Day 7 for cowbirds (n=4) (day after hunger-driven begging assay), focal nestlings were removed from the nest for the *in situ* muscle stimulation assays. The full procedure for stimulating the muscle can be found in the supplemental materials. Briefly, we implanted electrodes into the muscle through which we could send an electrical current to induce a muscle contraction, as well as a hook connected to a force transducer which measured the relative force produced by said contraction.

The aim of our study was to subject the **MC** to a stimulation mimicking the performance challenge of maintaining begging displays for longer than competitors. To emulate the constant contraction necessary to maintain this display performance, we stimulated the **MC** at 100 Hz to induce a single tetanic contraction [33]. This allowed us to investigate the rate and duration at which the muscle fatigued, which is observed and recorded using the progressive decay of tension (mV) measured by the force transducer that was attached to the contracting muscle. During the 60 second trial, the muscle slowly began to fatigue asymptotically after initial contraction; this could be observed by quantifying the rate at which force values returned to baseline (see traces in **Fig 2A**). Further, we investigated stimulation trains at 2 Hz and 5 Hz twitch kinetics (see Supplemental Methods; [30,32,34]) to measure the nestling’s ability to oscillate head and neck movements up to 2 times per second (biologically plausible; [35]) and 5 times per second (faster than observed head movements for begging in either of our tested species; Antonson – *personal observation*). All stimulation trains were performed in triplicate with at least 1 min in-between the end of one replicate and the start of another.

**Figure 2.**
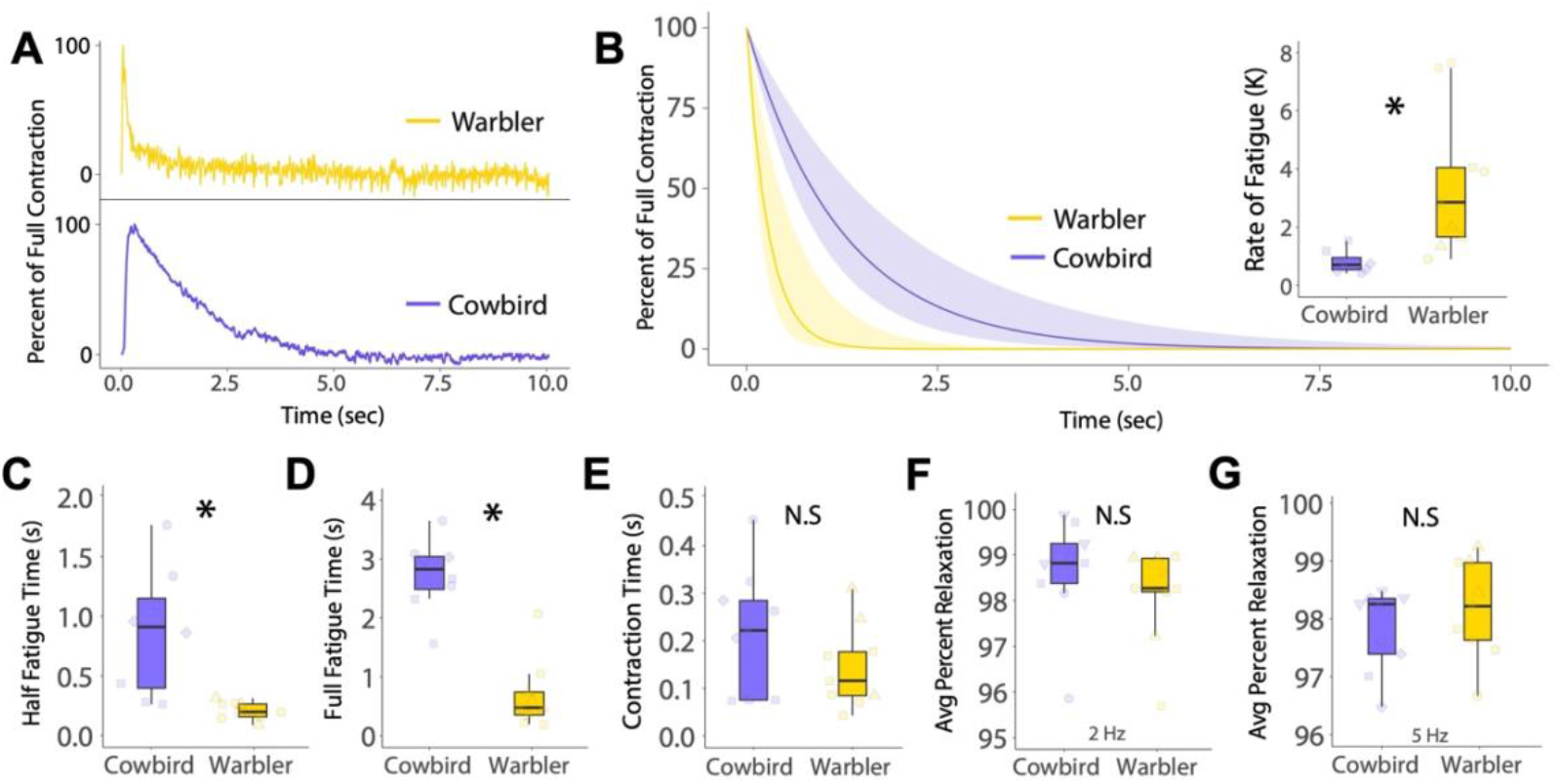
Response of the **MC** to *in situ* muscle performance assays. A) Force transducer traces in response to *in situ* stimulation (first 10 s shown). B) One-phase decay fatigue curves fitted to the decline in force from max contraction. Inset graph shows difference in fatigue rate constant (K) between species. C) Time to half muscle fatigue was significantly longer for cowbirds than their hosts, D) as was the time to full fatigue. E) Time to reach full contraction did not differ between species. Average percent muscle relaxation in response to low frequency twitch challenges at F) 2 Hz and G) 5 Hz were both not different between species. Box plots show interquartile-range and median. Asterisk (*) denotes significance (p<0.05), and N.S. denotes non-significant differences (p>0.05). Shapes represent n=3 cowbirds and n=3 warblers, with number of each shape showing replicates.

#### 3.2 Analysis of *in situ* fatigue challenge data

Complete muscle fatigue (0% of the maximum measured force) occurred when displacement on the force transducer was relieved to the point prior to stimulation (i.e., the signal returned to its baseline level). Baseline was calculated as the average value of force transducer displacement in the 0.5 seconds immediately preceding stimulation. Thus, when data values were between baseline (0%) and the point of max contraction (100%), the relative values of partial contractile force could be calculated by dividing the actual amount of contraction at a given point by the amount of contraction that would be necessary for full fusion relative to baseline. As such, we treated each 100 Hz stimulation as a continuous contraction, and measured the total time it took the muscle from the first stimulation to the point after maximal contraction (100%) where the relative contractile force first returned to baseline (0%). However, as the decline in force displacement was not linear and asymptotically approached 0%, we specifically measured the time it took force to fatigue to 50% (half fatigue time) and 10% (measure for full-fatigue time) of the maximal value. The goal of using these two measurements was to provide robust thresholds that would not be artificially inflated by the asymptotic nature of the data when comparing the fatigue-resistance of these two species. These data could be resolved for n=3 cowbirds and n=3 warblers.

Relative speeds of these contractions were measured as the time interval between the first stimulation pulse and the point of maximum relative contractile force. We likewise measured the relative force displacement measured by the force transducer for each contraction both to ensure no difference between species and to validate that the assay was equally sensitive across species (t=-1.031, p=0.361). Given this assay was not fully isometric and we did not measure length-tension relationships, this is not a direct measure of actual force produced.

### 4. Data Analysis

Data were analyzed using R (version 4.0.2), using an α = 0.05 for all models as the threshold for significant differences. We used R package ‘lme4’ to construct linear mixed effects models to test for the fixed effect species differences, with the random intercept of “*Individual”* accounting for replicate sampling and avoiding pseudoreplication. To determine the rate of fatigue during tetanic contraction, we fitted the decline in force from the maximum to a one-phase decay curve and calculated the rate constant (K) using the application Prism (version 10.1.0). All response variables were log-transformed to conform to normality more closely for statistical testing, while figures show raw values. To demonstrate the robustness of our results to restricted sample sizes, we also report standardized effect sizes (Cohen’s d) and bootstrapped 95% confidence interval estimates for said effect sizes for both the behavioral and performance models (**Fig S1**) as recommended in Nakagawa and Cuthill ([36]).

## Results

Begging displays between parasitic cowbirds and their warbler hosts are dynamic contests, which the more intensely begging cowbirds tend to win [17,24] (**Fig 1A**). Indeed, we found that cowbirds and warblers responded equally fast to provisioning stimuli, with both species taking ≈1 second to assume a full begging posture (Cohen’s d=-0.33, CI=-0.889, 0.179; t=-1.327, p=0.190; **Fig 1B**). However, cowbird nestlings performed significantly longer begging bouts than warbler nestlings (d=-1.53, CI = -2.79, -0.84; t=-6.817, p=0.001; **Fig 1C**). Similarly, cowbirds showed longer durations of total begging-associated activity compared with warblers (d=-0.927, CI=-1.725, - 0.517; t=-4.474, p=0.018; **Fig 1D**).

Next, we conducted a series of *in situ* muscle performance assays to examine whether behavioral differences in begging were attributed to divergent functional properties of the neck muscle that actuates this display. We first looked at endurance, testing how long the **MC** in cowbird and warbler nestlings could sustain a full contraction, much like those that are necessary to sustaining exaggerated begging bouts (**Fig 2A**). In this regard, the cowbird **MC** dramatically outperformed the warbler **MC**, with the cowbird muscle fatiguing at a significantly slower rate (d=0.789, CI=0.218, 1.622; t=2.845, p=0.046; **Fig 2B**). Specifically, the cowbird **MC** was able to maintain its contraction for a significantly longer duration than the warbler **MC**, before it reached both half-fatigue (d=-0.883, CI=-2.835, -0.267; t=-2.981, p=0.039; **Fig 2C**) and full-fatigue (d=-1.23, CI=-3.39, -0.27; t=-3.772, p=0.019; **Fig 2D**).

Meanwhile, we found that other **MC** performance attributes related to begging were statistically indistinguishable between the two species. For example, latency to reach maximum force in response to **MC** stimulation did not differ between cowbirds and warblers (d=-0.172, CI=-0.476, 0.004; t=-0.710; p=0.517; **Fig 2E**). Additionally, we assessed the **MC**’s twitch properties, and we found no difference in response when the muscles were stimulated at either 2Hz (d=-0.392, CI=-1.42, 0.186; t=-0.885, p=0.426) or 5Hz (d=0.254, CI=-0.148, 0.962; t=0.709, p=0.517) (**Fig 2F&G**). Taken together, these results point to clear species differences specifically in **MC** fatigue-resistance, but not other key performance attributes. These effects match our behavioral metrics, whereby superior fatigue-resistance likely confers the endurance that cowbirds need to support their exaggerated begging phenotype, relative to the shorter begging bouts of host warblers. To our knowledge, this is the first demonstration, to our knowledge, of a physiological performance trait that may underlie exaggeration in begging, or any other food-solicitation display.

## Discussion

Studies have previously hinted at the importance of muscle performance to support early-life success as a brood parasite [37,38]. Studies in cuckoos, for example, show that the **MC** has densely packed fiber composition to help chicks pip out of the thicker parasitic eggshell, which evolved to prevent egg damage by host parents [37]. Likewise, comparative work indicates that brood parasites perform greater rates of embryonic movement compared to host species [38]. Presumably, this serves to enhance *in ovo* development of muscular strength in parasites [38]. In parallel, we provide the first evidence that certain muscles in brood parasitic chicks are likely to influence the performance attributes of begging behavior and may be specialized for the task. This enhancement in muscular fatigue-resistance may serve as the mechanism by which cowbirds construct their parasitic niche, reducing host broods through a prolonged begging “war of attrition” [23].

The properties that may endow parasites with greater **MC** fatigue-resistance remain unclear. On one hand, it might be tempting to attribute species differences in MC endurance to effects of hatching asynchrony or differences in body size and developmental stage between cowbird and host chicks [24]. However, we doubt this is the case in this study for a variety of reasons. First, we find that cowbird and warbler chicks show similar MC performance features in response to *in situ* measures of twitch kinetics and time to reach maximum muscular contraction. If hatching asynchrony truly explained major differences in performance between these species, then we would have also likely expected these other measures to vary in kind. Second, prior behavioral work shows that cowbirds still beg longer than host chicks of red-winged blackbirds (*Agelaius phoeniceus*), which are similarly sized and hatch synchronously [39]. Cowbirds in this study also begged longer than larger brown thrashers (*Toxostoma rufum*) and for similar durations as smaller field sparrows (*Spizella pusilla*), both of which hatch a day later. Though these results should be interpreted in the context of satiation given that provisioning success between parasite and host differed for these two species (thrashers having more success than cowbirds and sparrows having less), while it did not when comparing cowbirds and blackbirds [39]. This provides evidence that innovations in extreme begging persist in cowbirds even in the absence of developmental lag and body size differences with the host. Accordingly, we suspect that species differences in begging endurance and fatigue-resistance arise from muscular specializations to actuate prolonged bouts of begging. For example simple changes in the expression profile of a handful of genes can lead to profound differences in muscular endurance properties [40–42]. Still, further work across multiple host species will likely be necessary to more finely determine whether the performance differences we report here are due to developmental factors or intrinsic species differences (or some combination of both).

From a broader evolutionary perspective, future work should test whether enhanced **MC** fatigue-resistance is a common feature across the six independently evolved instances of altricial avian brood parasitism [38]. We hypothesize that muscular adaptations to support exaggerated begging are likely shared with parasitic *Clamator* cuckoos [16] and potentially with *Vidua* finches [43],both of which are like cowbirds in that they do not directly kill host nestlings. The chicks of great spotted cuckoos (*Clamator glandarius*), for example, begs at a higher performance in both posture and duration compared to larger magpie host nestmates [16], and thus may do so by engaging similar specializations in muscular fatigue-resistance. Likewise, different muscular specializations outside of the **MC** may arise to facilitate the emergence of alternate extreme parasitic strategies for obligate brood parasitism such as chick killing, nestling ejection, etc. [44– 46].

Finally, our results highlight the importance of motor constraint to the evolutionary stability of behavioral strategies that help individuals negotiate familial conflict. Models exploring these dynamics focus primarily on costs of behavior, like begging, that limit the exaggeration of offspring need [1,47]. Our work, however, reveals that endurance imposes major limits to how individuals can accrue such costs [48]. Thus, evolution of behaviors to negotiate familial conflict should arise in response to a balance among not only behavioral costs and benefits, but also physiological constraints that limit performance. This perspective should be incorporated into the contemporary modeling that explores how and why parent-offspring conflict, as well as sibling rivalry, influences phenotypic evolution.

## Acknowledgements

We thank Nigel Anderson, Nicole Moody, Sofia Piggot, Lauren Renna, Rich Marsh, and Tom Roberts for helpful discussions about this paper.

## Supplementary Figures

**Figure S1.**
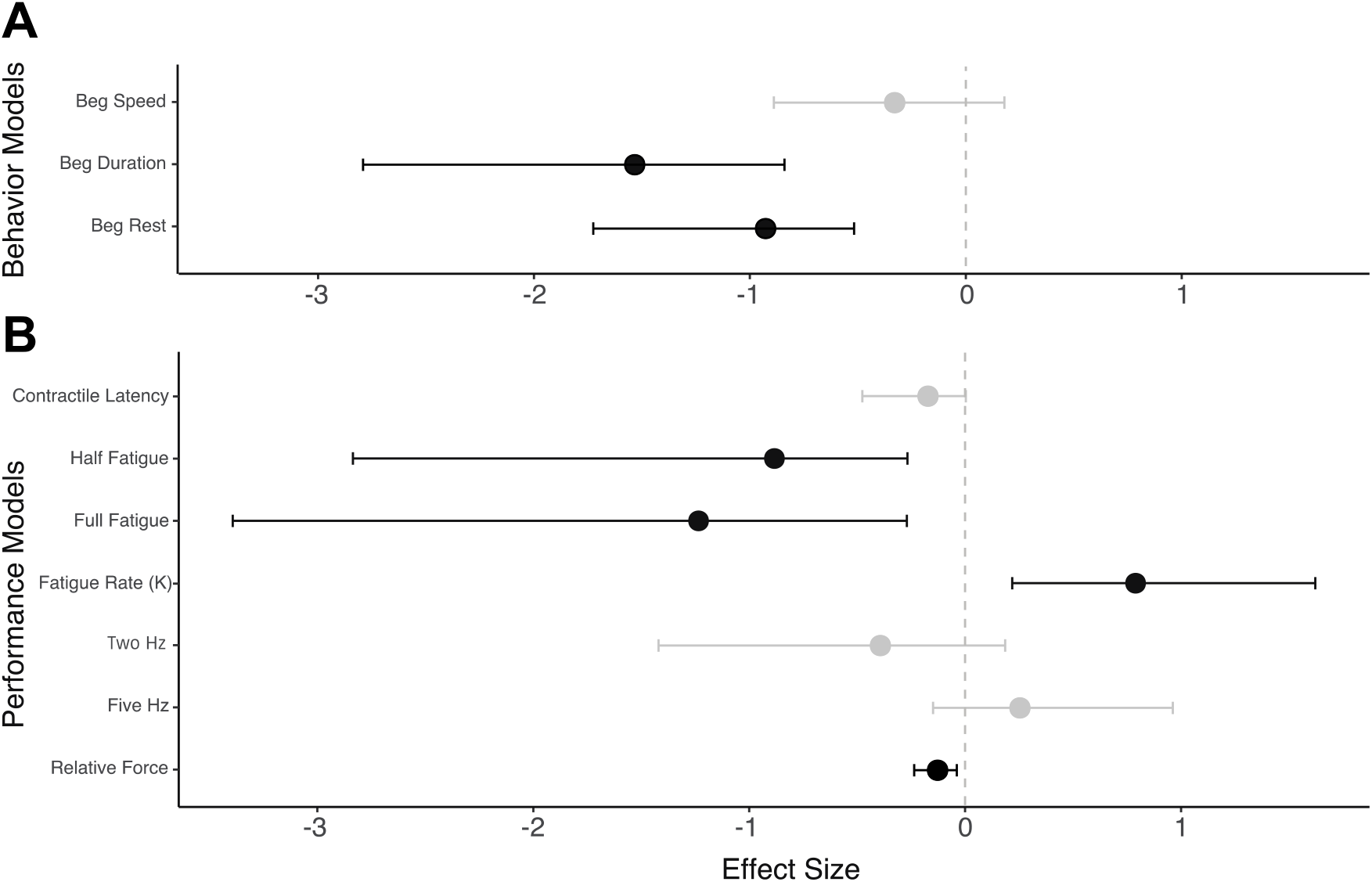
Cohen’s d standardized effect sizes calculated for models of A) behavioral comparisons and B) performance comparison between parasitic brown-headed cowbirds and host prothonotary warblers. 95% confidence intervals represented by whiskers around the mean effect size. Effect sizes in black were those where the confidence intervals did not overlap zero, suggesting that the data provide a robust estimate of the effect. Additionally, the effect sizes for all models reporting a significant difference (p<0.05) in the main text had substantial effect sizes, as an effect size of d=(±)0.8 is often considered a large effect.

