## supplemental materials for "Extreme fatigue-resistance in a brood parasite parallels a war of attrition in begging competition with hosts"

Supplementary Methods

*Quantification of hunger-driven begging assay times*

Time intervals for begging speed of response, endurance of begging bout, and total activity time were measured by analyzing the videos in Adobe Premiere (v24.0) Begging speed of response was measured as the time from the start of the stimulus to the time nestlings fully extended their neck upward in a peak begging posture. Endurance of the begging bout was the duration from the onset of movement in response to the stimulus, to the time point at which the neck retracted after extension during begging. Finally, total begging-associated activity was quantified as the total duration of movement after a stimulus until the nestling returned to rest by laying its head back on the nest substrate.

*In situ muscle stimulation procedure*

Birds were anesthetized with isoflurane (1-4% in $O_2$) and placed in surgical plane with the ventral side resting on the heating pad used to maintain body temperature. Once we confirmed the bird was anesthetized via toe pinch, we exposed the *musculus complexus* (**MC**) by a small nick (<1cm) in the skin atop the muscle. We implanted the the muscle with two small, insulated silver wire electrodes, which ran to a stimulator (AMS4100, A-M Systems, Sequim, WA, USA) and threaded a small stainless-steel hook into the muscle (0.1 mm diameter) between the positions of the implanted electrodes. This hook was connected by a monofilament line to a force transducer (Model FT03, Grass Technology, RI, USA), which was held taught by a weighted stand to measure relative force production by the contracting muscle in response to electrical stimulation. We immediately applied normal saline (0.9%) to the exposed muscle to prevent desiccation during our stimulation recordings, and we adjusted the placement of the force transducer to ensure that an appropriate level of tension was placed on the monofilament line to obtain high-quality muscle contractile and twitch recordings. Stimulation was performed with a current between 0.5-0.8mA to minimize body movement as done previously in other bird species of similar size (Fuxjager et al. 2016; Fuxjager et al. 2017; Miles et al. 2018). Further description of the experimental set up is available elsewhere (Fuxjager et al. 2016; Fuxjager et al. 2017; Miles et al. 2018).

Both the stimulator and force transducer were connected to a laptop through an A-D converter (Model NI USB-6212, National Instruments, TX, USA). The signal from the force transducer was amplified (5K–10K) and low-pass filtered (3000 Hz) using an AC/DC strain gage amplifier (Model P122, Grass Technologies). All recordings were collected using AviSoft (v4.2.22) and measured in Praat (v6.3.20). When the recordings were completed, the hook and electrodes were removed from the muscle, and the incision was closed first with silk suture and then sealed with Vetbond tissue adhesive. All procedures were completed at room temperature. Nestlings were returned to the nest and monitored daily until fledging with supplemental feeding during transport and at placement back in the nest. All experimental nestlings fledged after being returned to the nest.

*In situ approach considerations*

Invariably, due to the assay not being a fully isometric/lethal preparation of the muscle, small movements induced by stimulation or breathing had a minor effect on changing the resting position from before stimulation to after stimulation and thus shifted respective baseline values. For example, the body turning away from the force transducer would lead the new baseline to be below the original baseline, and the turning toward the force transducer would set the new baseline higher. We used three strategies to address this issue. 1) The force transducer was placed superior to the body while needing to be offset slightly to the right of the body to accommodate the anesthesia rig. 2) If baseline before stimulation was higher than the baseline after stimulation, we normalized the data using the baseline after stimulation to ensure the fatigue curves reached 100% fatigue after the full contraction that stimulation induced. Likewise, if the baseline after stimulation was higher, we normalized to the baseline prior to stimulation. By using this strategy, we conservatively applied the same effect despite directional shifting in body position by the birds. 3) By fitting one-phase decay curves to the data, we were able to effectively address the rate constant of fatigue independent of these small body movements, which provided us an additional, robust measurement for our trait of interest.

*Calculating percent relaxation for twitch kinetics*

To measure percent relaxation in the 2 Hz and 5 Hz treatments, we calculated baseline signal just prior to each trial, full relaxation for each trial, and percent relaxation after each individual stimulation pulse (as done previously in Fuxjager et al. 2016; Fuxjager et al. 2017; Miles et al. 2018; See Figure 2 in Fuxjager et al 2016 for specific details). Baseline signal was calculated as the average of 6 peaks and troughs immediately prior to stimulation within a given replicate. This allowed us to define partial relaxation as the difference between the maximum relaxation value between stimulation pulses and minimum relaxation divided by the value difference between 100% (maximum relaxation) and 0% (minimum relaxation, or full muscle fusion). Contractions were never fully tetanic between stimulation pulses for the twitch kinetic assay, so it was always possible to resolve percent relaxation. This data could be resolved in n=3 cowbirds and n=3 warblers.

*Validation of measures for consistency*

To ensure consistency across replicates (and evidence that the muscle fully recovered in the allotted time before further stimulation), we tested our performance parameters across replicates within each species. Rate of fatigue did not differ across replicates for cowbirds (t=0.406, p=0.704) or warblers (t=0.979, p=0.373). Time to full fatigue did not differ across replicates for cowbirds (t=-0.239, p=0.822) or warblers (t=-1.158, p=0.299), nor did half fatigue for cowbirds (t=1.331, p=0.253) or warblers (t=0.378, p=0.717). Force production was also indistinguishable across replicates for both cowbirds (t=-2.127, p=0.100) and warblers (t=-1.443, p=0.209). Finally, in response to 2 Hz twitch kinetics, there was also no difference across replicates for cowbirds (t=-0.362, p=0.732) or warblers (t=-0.378, p=0.716), nor at 5 Hz for cowbirds (t=-0.092, p=0.930) or warblers (t=-0.202, p=0.848).

**Supplementary Figures**

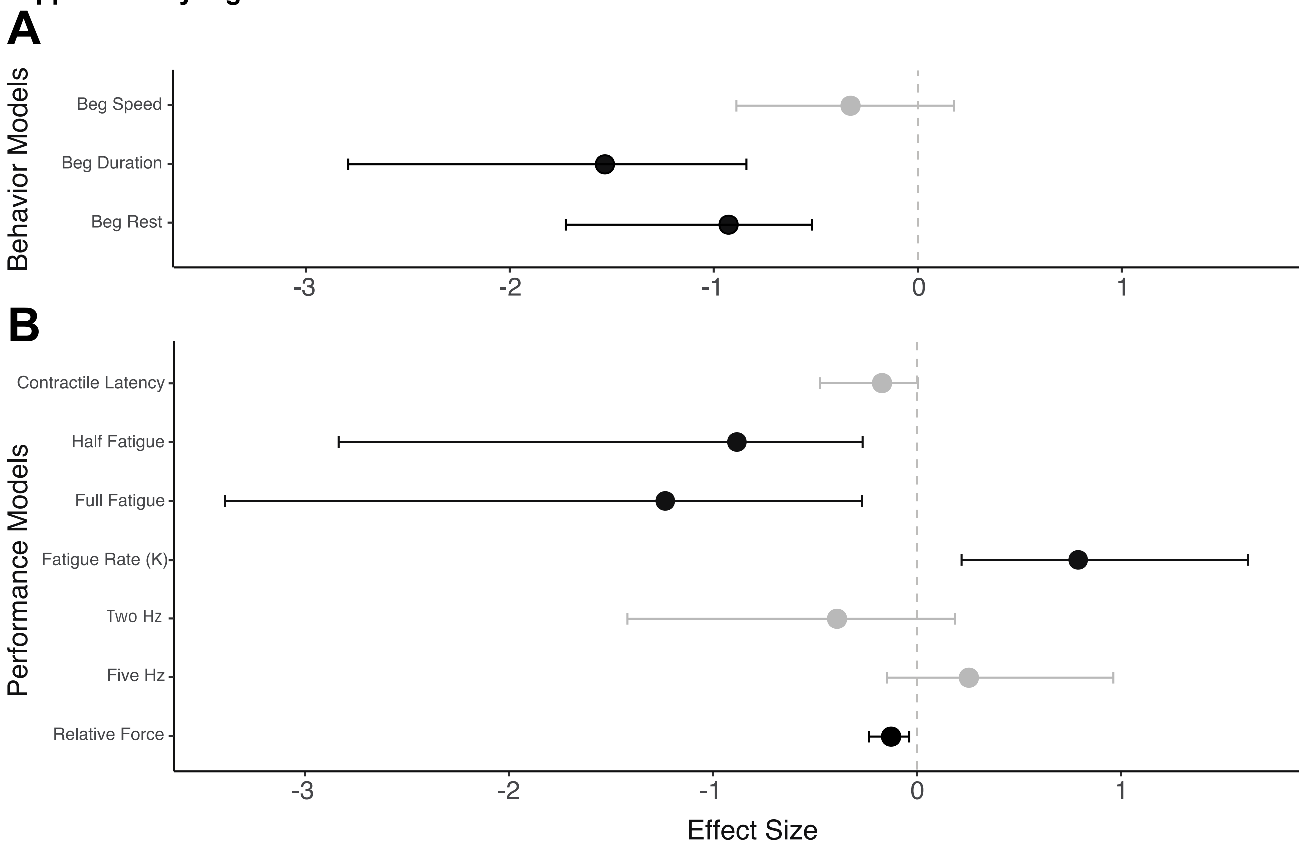


**Figure S1.** Cohen's d standardized effect sizes calculated for models of A) behavioral comparisons and B) performance comparison between parasitic brown-headed cowbirds and host prothonotary warblers. 95% confidence intervals represented by whiskers around the mean effect size. Effect sizes in black were those where the confidence intervals did not overlap zero, suggesting that the data provide a robust estimate of the effect. Additionally, the effect sizes for all models reporting a significant difference ($p<0.05$) in the main text had substantial effect sizes, as an effect size of $d=(\pm)0.8$ is often considered a large effect.
